# Beta rhythm events predict corticospinal motor output

**DOI:** 10.1101/588632

**Authors:** Sara J Hussain, Leonardo G Cohen, Marlene Bönstrup

**Affiliations:** Human Cortical Physiology and Neurorehabilitation Section, National Institute of Neurological Disorders and Stroke, National Institutes of Health. Bethesda, MD, 20892.

**Author notes:** **Corresponding author:** Sara J Hussain, PhD 10 Center Dr. Rm 7D49 Bethesda, MD, USA. **Author contributions:** SJH and MB conceived of and designed research; SJH collected data, SJH and MB analyzed data; SJH, LGC and MB wrote the manuscript. **Competing interests:**The authors declare no competing interests.

## Abstract

The beta rhythm (15-30 Hz) is a prominent signal of sensorimotor cortical activity. This rhythm is not sustained but occurs non-rhythmically as brief events of a few (1–2) oscillatory cycles. Recent work on the relationship between these events and sensorimotor performance suggests that they are the biologically relevant elements of the beta rhythm. However, the influence of these events on corticospinal excitability, a mechanism through which the primary motor cortex controls motor output, is unknown. Here, we addressed this question by evaluating relationships between beta event characteristics and corticospinal excitability in healthy adults. Results show that the number, amplitude, and timing of beta events preceding transcranial magnetic stimulation (TMS) each significantly predict MEP amplitudes. However, beta event characteristics did not explain additional MEP amplitude variance beyond that explained by mean beta power alone, suggesting that conventional beta power measures and beta event characteristics similarly capture natural variation in human corticospinal excitability. Despite this lack of additional explained variance, these results provide first evidence that endogenous beta oscillatory events shape human corticospinal excitability.

## Introduction

The primary motor cortex influences motor output through a balance between excitation and inhibition within cortical microcircuits that synapse onto layer V corticospinal neurons. The excitability of these circuits can be measured non-invasively using transcranial magnetic stimulation (TMS), which elicits motor-evoked potentials (MEPs; 1). Excitability of these circuits is not static but instead fluctuates spontaneously and is in part determined by the phase and power of endogenous sensorimotor oscillatory activity (2–5).

We recently reported that sensorimotor beta activity positively relates to corticospinal excitability, with higher beta power predicting larger MEP amplitudes (5). However, the beta rhythm is not sustained but emerges as brief episodes that typically last between 1-2 oscillatory cycles, termed beta events. Biophysical modeling of cortical microcircuits suggests that beta events emerge from interactions between a broad, excitatory synaptic drive to proximal dendrites of pyramidal neurons and a shorter, stronger excitatory drive to distal dendrites of pyramidal neurons (6,7). Further, characteristics of beta events, including their number, duration, amplitude and timing with respect to stimulus presentation (Figure 1), are associated with variability in somatosensory perception (8) and motor performance (9). However, the relationship between beta events and corticospinal excitability, a basic mechanism by which the primary motor cortex shapes motor output (10), is unknown.

**Figure 1.**
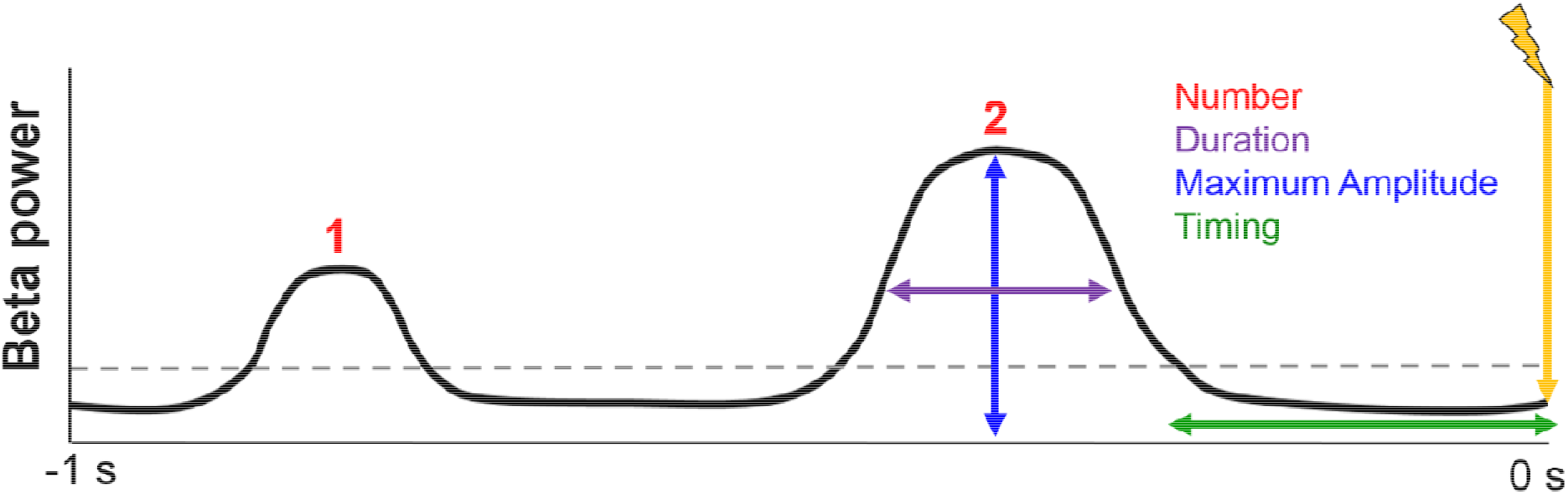
Illustration of beta event characteristics. Beta events were defined as portions of the beta band timeseries data exceeding one of two thresholds (75^th^ or 82^nd^ percentile power) for at least 1.5 cycles. Events are characterized by their number (red), duration, (purple), maximum amplitude (blue), and timing (green) relative to TMS delivery (yellow). See *Methods* for details on event characteristic calculation.

Here, we evaluated the relationship between corticospinal excitability and various characteristics of beta events using simultaneous TMS and EEG recorded from healthy volunteers (5). We report that several beta event characteristics significantly predict MEP amplitudes, but that these characteristics do not explain additional variability in MEP amplitudes compared to that explained by mean beta power alone. Overall, findings support the biological relevance of multiple beta event characteristics while also demonstrating that these characteristics do not outperform conventional beta power measurements when predicting human corticospinal excitability.

## Results

### Beta event characteristics

We first identified beta events from EEG data recorded over the sensorimotor cortex during TMS delivery. Beta events were identified as portions of beta band timeseries data that exceeded a given percentile power threshold within individuals. To evaluate reproducibility of our findings across different thresholds, we used both literature-based (75^th^ percentile, 9) and empirically-defined (82^nd^ percentile, 8) thresholds. Beta events were defined as time periods during which mean beta power exceeded these thresholds, were at least 1.5 cycles long (9) and fell within 1 second before each TMS pulse. The frequency band of the beta time series was defined as the individual peak frequency within the beta band (15-30 Hz, Figure 2A). Multiple morphological characteristics were then extracted from these beta events for each pre-stimulus period, including beta event duration, maximum amplitude, and timing of the last event relative to the TMS pulse (Figure 1).

**Figure 2.**
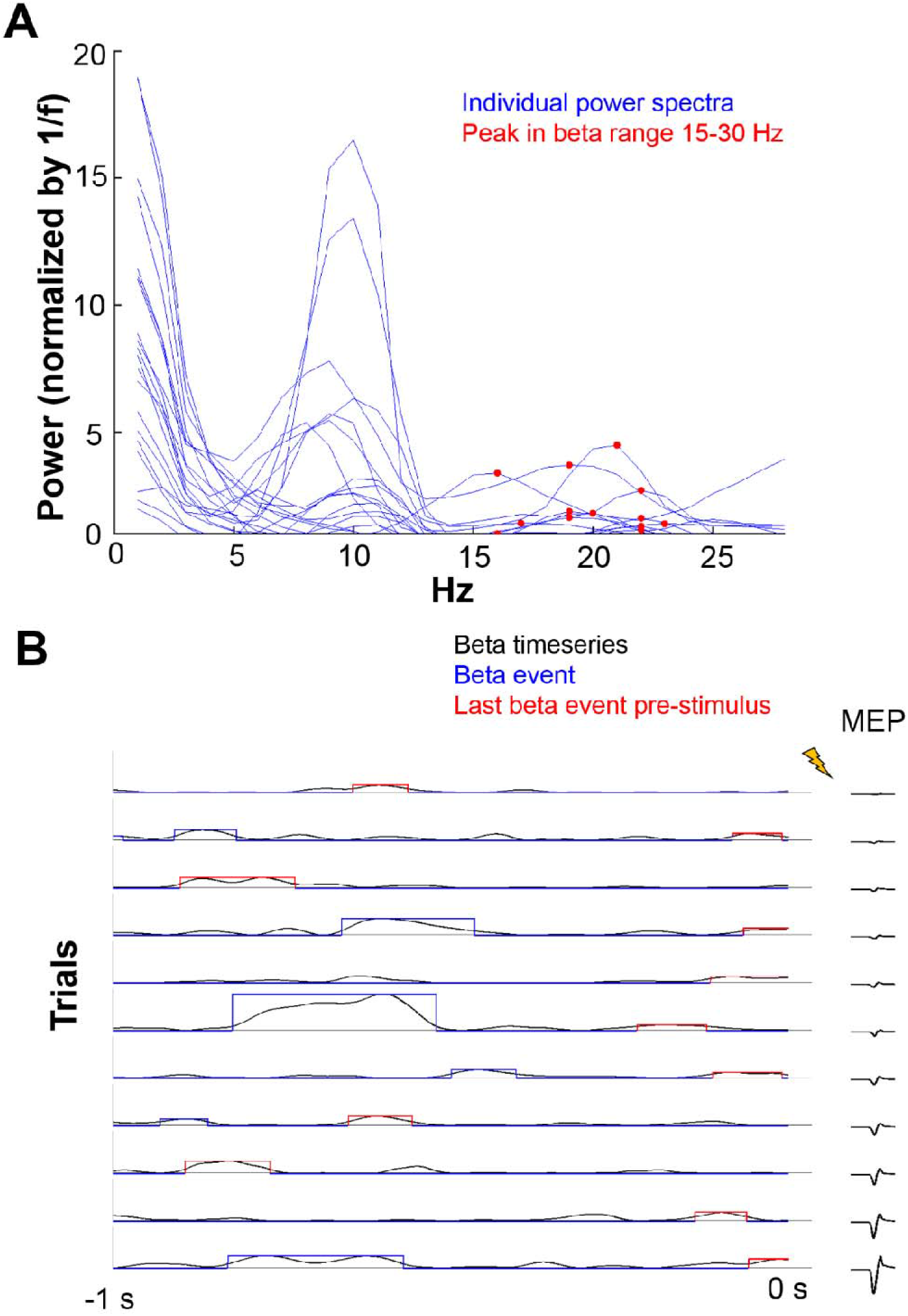
Beta power timeseries, beta events and event characteristics. A) Power spectral densities for each subject (blue lines) and individual peak beta power frequencies between 15-30 Hz (red dots). B) 10 trials from one representative subject are depicted, arranged according to MEP size. Beta timeseries data are shown in black. Events are indicated in blue and red, with their timing, duration and maximum amplitude given by the location, width, and height of the rectangular shape around the beta timeseries, respectively. The last event before the TMS pulse, for which maximum amplitude and duration were used for statistical modeling, is marked in red. Corresponding MEP traces are depicted in black to the right of each beta timeseries.

As previously described, the sensorimotor beta rhythm was comprised of transient, non-rhythmic burst-like events of high beta power. Events occurred at a rate of 1.24 per second (1.17 ± 0.016 [SEM] events per 0.95 s pre-stimulus period) and lasted for ~110-120 ms (114 ± 0.9 [SEM] ms). Further, beta event characteristics were moderately yet significantly co-linear with mean beta power (Spearman’s correlation per subject, number [mean R=0.58 ± 0.02], duration [R=0.26 ± 0.04], and amplitude [R=0.54 ± 0.02], p<0.05 for all]).

### Beta event characteristics predict MEP amplitude

We then used linear mixed-effects modeling to evaluate whether the presence of a beta event in a given pre-stimulus period influenced MEP amplitude, and if so, which specific beta event characteristics were related to MEP amplitudes. This approach demonstrated that the presence of a beta event during the pre-stimulus period was associated with larger MEP amplitudes (model fit on all trials: N=8237, model estimate=0.08, p=6.80^−5^). Further, multiple beta event characteristics were significantly associated with MEP amplitudes (Table 1), including the number of beta events within the pre-stimulus period (Figure 3A), beta event maximum amplitude (Figure 3B), and timing of the last event relative to TMS delivery (Figure 3C). Positive model estimates for beta event number and maximum amplitude indicate that larger MEP amplitudes were associated with a greater number and larger events (Figures 3A, 3B). In contrast, the negative model estimate for beta event timing indicates that larger MEP amplitudes were associated with events that occurred more closely in time to TMS delivery (Figure 3C). 15 of 20 subjects showed a positive relationship between beta event number and MEP amplitudes, 16 of 20 subjects showed a positive relationship between beta event maximum amplitude and MEP amplitudes, and 14 of 20 subjects showed a negative relationship between beta event timing and MEP amplitudes. Results were similar regardless of the event detection threshold used (Table 1).

**Table 1.** Relationships between MEP amplitude and beta metrics across two different event-definition thresholds. Asterisks (*) indicate significance at an alpha of 0.05. Note that cross-validated rho and normalized RMSE values are very similar regardless of the beta metric and/or event threshold used.

|  | Beta metric | Model estimate | Model estimate p-value | AIC | Cross-validated rho | Cross-validated RMSE |
| --- | --- | --- | --- | --- | --- | --- |
| 75<sup>th</sup> %ile threshold<br>69% event trials<br>N=5663 | Event number | 0.05 | 4.7<sup>-7*</sup> | 21428 | 0.62* | 0.13 |
|  | Event duration | 0.31 | 9.3<sup>-2</sup> | 14581 | 0.62* | 0.13 |
|  | Event amplitude | 0.01 | 2.3<sup>-3*</sup> | 14575 | 0.62* | 0.13 |
|  | Event timing | -0.10 | 2.9<sup>-2*</sup> | 14579 | 0.62* | 0.13 |
|  | Mean beta power | 0.07 | 1.8<sup>-6*</sup> | 14561 | 0.62* | 0.13 |
| 82<sup>nd</sup> %ile threshold<br>54% event trials<br>N=4477 | Event number | 0.06 | 6.0<sup>-8*</sup> | 21424 | 0.61* | 0.13 |
|  | Event duration | 0.20 | 3.9<sup>-1</sup> | 11597 | 0.62* | 0.13 |
|  | Event amplitude | 0.01 | 1.4<sup>-3*</sup> | 11587 | 0.62* | 0.13 |
|  | Event timing | -0.09 | 7.4<sup>-2</sup> | 11594 | 0.62* | 0.13 |
|  | Mean beta power | 0.07 | 8.0<sup>-6*</sup> | 11577 | 0.62* | 0.13 |
| All trials<br>N=8327 | Mean beta power | 0.08 | 4.6<sup>-11*</sup> | 21410 | 0.61* | 0.13 |

**Figure 3.**
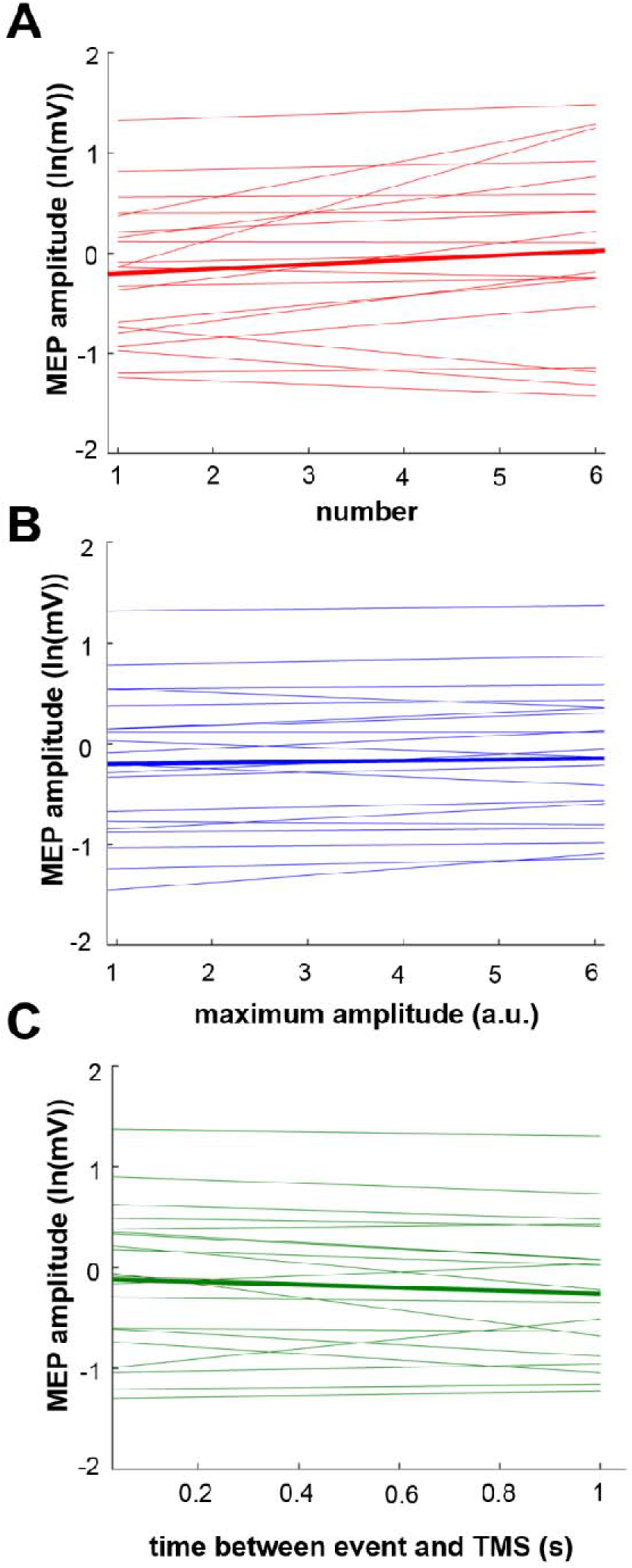
Beta event characteristics predict MEP amplitude. Beta event number and maximum amplitude were both positively related to MEP amplitude, indicating that a greater number of events, as well as larger events, preceded larger MEPs. The timing of the last event relative to TMS delivery was negatively associated with MEP amplitude, indicating that MEPs were larger when an event occurred more closely in time to TMS delivery. Thick lines indicate group-level model fits and thin lines indicate individual subject regression lines (fit using least-squares method).

**Figure 4.**
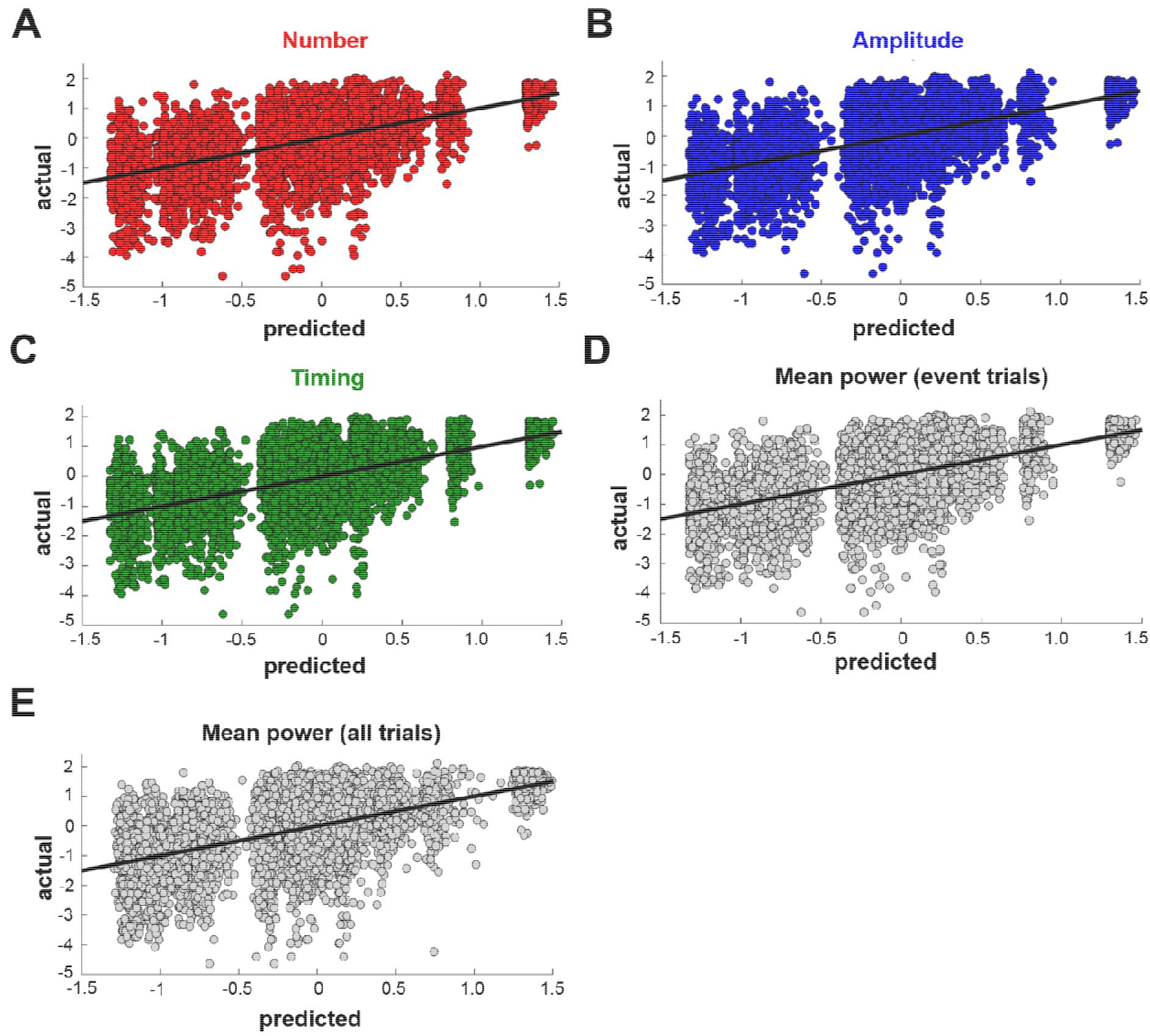
Prediction of MEP amplitudes based on beta event metrics. K-fold (k=10) cross-validation was performed to evaluate the ability of different beta event metrics (including event number [A], amplitude [B], timing [C], as well as mean beta power [D-E]) to predict MEP amplitudes. Actual MEP amplitudes (y-axis) are plotted against predicted (x-axis). All plotted data were obtained using the 75^th^ percentile threshold. For all metrics, cross-validated rho values and normalized RMSE values indicated moderate predictive performance (rho values=0.61-0.62, normalized RMSE=0.13).

After establishing that beta event characteristics were significantly related to MEP amplitudes, we then determined the ability of beta event characteristics to predict MEP amplitudes using cross-validation (11). We implemented 10-fold cross-validation for each linear mixed-effect model (Table 1), using Spearman’s correlation and root-mean square error (RMSE) to evaluate relative and absolute predictive performance, respectively. Cross validation revealed that both the relative and absolute prediction of MEP amplitudes by beta event characteristics was moderate (mean Spearman’s rho across folds = 0.61-0.62; p<0.001 for each fold [relative prediction]; Normalized RMSE = 0.13 [absolute prediction], Table 1). Linear relationships indicating relative prediction of MEP amplitudes by beta event characteristics are depicted in Figure 4. Thus, predictive models utilizing beta event characteristics and mean beta power predicted spontaneous fluctuations in MEP amplitudes to similar extents. Results were again consistent across the two event detection thresholds used (Table 1).

### Beta event characteristics have no additional predictive value beyond mean beta power

Given the only moderate co-linearity between beta event characteristics and mean beta power, it is possible that beta event characteristics could explain additional variance in MEP amplitude that is not accounted for by mean beta power. To test this, we performed likelihood-ratio tests between simpler models containing only mean beta power as a fixed effect and more complex models containing both mean beta power and each beta event characteristic of interest (either number, duration, maximum amplitude, or timing) as fixed effects. Likelihood-ratio tests consistently revealed that the more complex models containing two fixed effects (i.e., mean beta power and a given beta event characteristic) were not superior to simpler models containing only mean beta power as fixed effects (number, p=0.17; amplitude, p=0.96; timing, p=0.18; duration, p=0.89). Results were again similar regardless of the event detection threshold used.

## Discussion

In this study, we evaluated the influence of non-rhythmically occurring brief events of strong beta activity, likely the biologically relevant elements of the beta rhythm (7–9), on corticospinal excitability assessed with single-pulse TMS. Here, we report that numerous, larger, and recent beta events predict larger MEP amplitudes. Consistent with our previous work (5), higher mean beta power was also associated with larger MEP amplitudes. However, the ability of beta event characteristics and mean beta power to predict MEP amplitudes was very similar, and a quantitative model-building approach revealed that beta event characteristics did not explain any additional variance in MEP amplitudes beyond what could be accounted for by mean beta power alone.

The beta rhythm is closely tied to activity within the sensorimotor system, as it specifically reflects voluntary motor behavior (including movement imagery, preparation, and execution; 12-14) and somatosensory performance (8,15,16). Coherence between cortical oscillatory activity and peripheral muscle activity is most prominently observed in the beta range (17,18), with this coherence being causally relevant for corticospinal signal transmission in healthy adults (19). In Parkinson’s disease, exaggerated beta activity is observed in the subthalamic nucleus and cortex (20,21), and the therapeutic effects of dopaminergic treatment and deep brain stimulation have been proposed to occur through elimination of this pathological beta activity (22,23). Previously, we and others have reported that pre-stimulus beta power correlates with corticospinal excitability in the resting brain (5,24,25), and a recent study also demonstrated that applying beta transcranial alternating current stimulation (tACS) to the motor cortex increases corticospinal excitability and beta power (26). Nonetheless, these studies have produced mixed findings, with some reporting positive relationships between corticospinal excitability and MEP amplitude (5,26) and others reporting negative relationships (24,25). These seemingly contradictory results may however be caused by different transcranial magnetic stimulation intensities used across studies, leading to activation of different neural populations (27,28). For example, lower intensity stimulation activates descending corticospinal neurons via interneurons in layers II/III, indirectly evoking descending corticospinal volleys (I-waves; 27,29). In contrast, higher intensity stimulation indirectly evokes descending volleys and can also directly evoke descending corticospinal volleys by activating axons of layer V corticospinal neurons themselves (I-and D-waves, respectively; 27,29). It is therefore conceivable that the direction of the relationship between beta activity and corticospinal excitability crucially depends on whether D-waves are evoked via direct activation of layer V corticospinal neurons. Our results combined with these other reports (24–26) suggest that beta activity may reflect inhibition of layer II/III interneurons and simultaneous excitation of layer V corticospinal neurons.

Beta event characteristics, including their number, amplitude, and duration, all contribute to mean beta power measurements, are significantly co-linear and are therefore conceptually and analytically related. As in the current study, previous work reported relationships between beta event characteristics and sensorimotor performance (8,9) but did not demonstrate that beta event characteristics could explain additional variance in sensorimotor performance compared to mean beta power alone. Consistent with this work, our results show that beta event characteristics and mean beta power predict MEP amplitudes to a similar extent, with beta event characteristics failing to add predictive value beyond mean beta power.

Despite similar predictive performance, measuring beta events and analyzing their waveforms enables a deeper understanding of specific cellular and circuit-level neural generators via simulation and interpretation of experimental observations (30) which is a step forward from mere quantification of oscillatory power. Of note, Sherman and colleagues (6) recently proposed a biophysical model that accounts for the specific shape of beta event waveforms across species and recording modalities. According to this model, beta events arise from simultaneous weak proximal and strong distal excitatory drives to pyramidal neurons in layers II/III and V. This same excitatory projection likely also synapses on inhibitory neurons in superficial cortical layers (6,31,8), which is consistent with the notion that sensorimotor beta activity produces an inhibitory cortical state (32,33). Low-intensity TMS is known to preferentially activate layer V corticospinal neurons trans-synaptically through layer II/III excitatory and inhibitory interneurons, as well as cortico-cortical connections (34,35). This indirect activation of layer V corticospinal neurons produces I-waves at the spinal level and MEPs at the muscle level. During low-intensity TMS, MEP amplitude variation therefore depends on the excitability state of the cortical interneuronal pool. During beta events, the excitatory drive to inhibitory interneurons in superficial cortical layers may thus increase the inhibitory tone of this interneuron pool, generating an inverse relationship between beta event presence and the elicited MEP size as previously reported (24,25). High-intensity TMS, in contrast, also directly activates the axons of layer V corticospinal neurons (34,35), which could feasibly overcome any inhibitory tone produced by inhibitory interneurons in superficial cortical layers. The strong excitatory drive to layer V corticospinal neurons that generates beta events may amplify this direct axonal activation by increasing the pool of layer V corticospinal neurons that are recruited by TMS, producing a positive relationship between beta event presence and MEP size as observed here. Thus, even when beta events do not predict an experimental outcome more strongly than mean beta power (8,9, and the current work), accounting for event characteristics can offer a more physiologically-informed framework for interpreting empirical findings.

Despite the conceptual advantages of beta event characteristics, it is important to note that there are some scenarios in which measuring mean beta power is preferable to measuring beta event characteristics. These include real-time applications, like brain state-dependent stimulation interventions (3,36) and brain-computer interfaces (37,38) in which the time needed to accurately estimate beta activity is crucial, or experimental designs in which only very short data segments that do not allow reliable extraction of beta timeseries are available. Our findings suggest that in these situations, mean beta power is suitable for quantifying the strength of the beta rhythm. Further, several of the models tested here did not show an equal predictive performance across the full range of observed MEP amplitudes, indicated by gaps across the predicted MEP sizes in Figure 4. These gaps were more prominent when only trials containing beta events were used, indicating that models incorporating mean beta power from all trials are likely to perform better in scenarios where continuous predictions of MEP amplitude is desired. In contrast, models incorporating either beta event characteristics or mean beta power may perform equally well in scenarios when it is only necessary to determine if an upcoming MEP amplitude will be large or small. Despite quantitatively similar predictive performance across all beta metrics tested here, the precise beta metric that is most appropriate for a given scenario depends qualitatively on experimental goals.

In summary, we demonstrate that the presence of beta events and their specific characteristics predict MEP amplitude, but that the ability of beta events to predict MEP amplitude does not exceed that of that of mean beta power. Specifically, beta event characteristics did not explain any additional variance in MEP amplitudes compared to mean beta power, indicating that these different measures are highly conceptually related. Despite this conceptual and predictive similarity, we suggest that defining beta event characteristics offers multiple advantages, including a more precise understanding of the beta rhythm’s temporal dynamics and a more physiologically-grounded framework for interpreting experimental results. Overall, our findings reinforce the importance of the beta rhythm in human motor control by documenting its role in determining corticospinal excitability, a mechanism through which the primary motor cortex controls human motor output.

## Methods

### Participants

Healthy adults (N=20, 6 F, 14 M, age = 30 ± 1.59 years) participated in this study, which was approved by the National Institutes of Health Combined Neuroscience Section Institutional Review Board. All subjects provided their written informed consent before participation. All study procedures were performed in accordance with local regulations for human subjects’ research. Data acquired for this study have been published previously (5).

### Transcranial magnetic stimulation (TMS)

The scalp hotspot for the left first dorsal interosseous muscle (L. FDI) was first identified over the hand representation of the right motor cortex using single-pulse monophasic TMS (MagStim 200^2^, MagStim Co. Ltd, UK) using a figure-of-eight coil held at ~45° relative to the mid-sagittal line, corresponding to a posterior-to-anterior current direction across the central sulcus (39). Resting motor threshold (RMT) was then determined using an automatic threshold-tracking algorithm (adaptive PEST procedure; 40). After hotspot identification and thresholding, 600 single, monophasic open-loop TMS pulses were delivered to the scalp hotspot for the L. FDI at 120% of RMT (inter-pulse interval: 5 s with 15% jitter). This intensity was chosen to elicit a motor-evoked potential (MEP) on each trial. During TMS, EEG recordings were simultaneous obtained and coil position was monitored online using frameless neuronavigation (BrainSight, Rogue Research, Montreal).

### EMG processing

MEP amplitudes resulting from TMS delivery were calculated offline from EMG signals. Peak-to-peak MEP amplitudes were defined as the difference between maximum and minimum voltage deflections within +0.020 to +0.040 s after each TMS pulse. We also calculated EMG signal power within −0.025 to −0.005 s before each pulse. Trials in which more than half of these pre-stimulus samples exceeded an empirically-defined upper limit (75^th^ percentile + 3*InterQuartile Range of EMG power) were identified and excluded due to excessive voluntary motor activity. In addition, trials in which MEPs showed low correlations (r ≤ 0.40) with the mean MEP signal, reflecting further contamination by voluntary motor activity, were also excluded. Resulting MEP amplitudes were then ln-transformed to reduce skew (5).

### EEG processing

EEG data were segmented offline into 6 s intervals (−3 and +3 s relative to each TMS pulse, with TMS delivery occurring at 0 s) and re-referenced to the average reference. Sensorimotor rhythms were extracted from the right motor cortex by applying the Hjorth transform of the electrode closest to the contralateral motor cortex (central = C4, surround = FC2, FC6, CP2, CP6; 4,5,41). Intervals containing artifacts within 1 second preceding each TMS pulse were automatically identified (maximum amplitude deflections of > 50 μV and kurtosis > 4) and excluded from analysis. After excluding intervals contaminated by voluntary EMG activity and/or EEG artifacts, 64 ± 10% (mean ± SD) of all intervals remained.

Pre-stimulus data were spectrally decomposed into power spectral density estimates between 1 and 30 Hz (Fast Fourier Transform, with Slepian sequences as tapers, smoothing frequency=2 Hz, [6,8]). Each subject’s individual beta frequency (IBF) was visually identified as the frequency between 15 and 30 Hz at which spectral power was maximal. Time-resolved estimates of beta power at each subject’s IBF were obtained using a 5-cycle wavelet decomposition. To avoid edge effects, the pre-stimulus timeseries (3 s) was mirrored at the time of the TMS pulse to create a 6 s long segment prior to wavelet decomposition. Resulting beta band timeseries were individually normalized by dividing by the mean beta power during the pre-stimulus period. Trials were then defined as the pre-stimulus portions of the beta band timeseries, covering a time window of −1 to −0.05 s before TMS delivery.

We identified beta events as portions of the beta band timeseries data that exceeded a given percentile power threshold within individuals. To evaluate reproducibility of our findings across different thresholds, we used two different approaches. First, we chose the 75^th^ percentile as the first threshold (based on previously published work; 9). Second, we empirically determined a threshold by correlating the percentage of time samples within beta events in the pre-stimulus period against the mean beta power of the pre-stimulus period. This was done over a range of possible thresholds (range=50-98). We then chose the percentile at which the group mean correlation (Pearson’s) curve was maximal (8) as our second beta event threshold, which corresponded to the 82^nd^ percentile. Based on these two thresholds (75^th^ and 82^nd^ percentiles), beta events were defined as time periods during which the mean beta power exceeded these thresholds, were at least 1.5 cycles long (9) and fell within the trial.

We extracted multiple morphological characteristics from the identified beta events for each trial (8). First, the number of events were counted. Then, for each event, we calculated the duration (defined as the time during which event power was greater than half of the event’s maximum power) and maximum amplitude (event maximum amplitude). For each trial, the timing of the last event prior to the TMS pulse was also determined. High amplitude segments at trial edges were considered events if the maximum power was reached within the timeseries (i.e. power was declining at the trial’s edge). The duration of these edge events was estimated as twice the pre-maximum amplitude duration. In the case of several events per trial, the maximum amplitude and duration of the last event prior to the TMS pulse were used for statistical testing.

### Statistical analysis

Linear mixed-effects modeling of trial-by-trial relationships between beta rhythm metrics and MEP amplitudes was performed (fixed effects of METRIC [number, duration, maximum amplitude, timing, or mean power], random intercepts of SUBJECT). Separate models were fit for each beta rhythm metric and the model estimates, p-values and Akaike Information Criterion (AIC) were determined. The additional predictive value of any beta event characteristics beyond mean power was then assessed by fitting linear mixed-effects models with fixed effects of mean power and the event characteristic of interest (fixed effects of MEAN POWER and CHARACTERISTIC [either number, duration, maximum amplitude, or timing]). Comparisons between models containing the single fixed effect of mean power versus the model containing two fixed effects of mean power and the event characteristic of interest were performed using likelihood ratio tests.

To test the generalizability of each model on independent data, we performed k-fold cross-validation (k=10). For each fold, data were divided into a training set (90% of available trials per subject) and a test set (10% of available trials per subject). Linear mixed-effects models for the predictor of interest (number, duration, maximum amplitude, timing, or mean power) were then fit to the training set, and the resulting model was used to predict MEP amplitude for each trial of the test set. The relative and absolute predictive performance of each model was quantified using Spearman’s correlation between predicted and actual MEP amplitudes (relative predictive performance; 1 indicating perfect relative prediction; 42) and the root-mean square error (RMSE) between predicted and actual MEP amplitudes (absolute predictive performance; 0 indicating perfect absolute prediction). After all folds were complete, correlation coefficients and RMSE values were averaged across folds. Because RMSE measures depend on the units of the dependent variable, we then normalized the averaged RMSE value to the range (maximum – minimum) of observed ln-transformed MEP amplitudes for interpretability.

## Data availability statement

The datasets analyzed during the current study are available from the corresponding author on reasonable request.

## Acknowledgements

This work was supported by the Intramural Research Program of the National Institute of Neurological Disorders and Stroke. SJH is supported by an NINDS Intramural Competitive Fellowship. MB is supported by the German National Academy of Sciences Leopoldina (Fellowship Program grant number LPDS 2016-01). The authors would like to thank Lutz Krawinkel for useful ideas for data analysis.

